# Structural Elucidation of Intact Rough-Type Lipopolysaccharides using Field Asymmetric Ion Mobility Spectrometry and Kendrick Mass Defect Plots

**DOI:** 10.1101/2023.06.21.545950

**Authors:** Abanoub Mikhael, Darryl Hardie, Derek Smith, Helena Pětrošová, Robert K. Ernst, David R. Goodlett

## Abstract

Lipopolysaccharide (LPS) is a hallmark virulence factor of Gram-negative bacteria. It is a complex, structurally heterogeneous mixture due to variations in number, type, and position of its simplest units: fatty acids and monosaccharides. Thus, LPS structural characterization by traditional mass spectrometry (MS) methods is challenging. Here, we describe the benefits of field asymmetric ion mobility spectrometry (FAIMS) for analysis of intact R-type lipopolysaccharide complex mixture (lipooligosaccharide; LOS). Structural characterization was performed using *Escherichia coli* J5 (Rc mutant) LOS, a TLR4 agonist widely used in glycoconjugate vaccine research. FAIMS gas phase fractionation improved the (S/N) ratio and number of detected LOS species. Additionally, FAIMS allowed the separation of overlapping isobars facilitating their tandem MS characterization and unequivocal structural assignments. In addition to FAIMS gas phase fractionation benefits, extra sorting of the structurally related LOS molecules was further accomplished using Kendrick mass defect (KMD) plots. Notably, a custom KMD base unit of [Na-H] created a highly organized KMD plot that allowed identification of interesting and novel structural differences across the different LOS ion families; i.e., ions with different acylation degrees, oligosaccharides composition, and chemical modifications. Defining the composition of a single LOS ion by tandem MS along with the organized KMD plot structural network was sufficient to deduce the composition of 179 LOS species out of 321 species present in the mixture. The combination of FAIMS and KMD plots allowed in-depth characterization of the complex LOS mixture and uncovered a wealth of novel information about its structural variations.

## 1. Introduction

Lipopolysaccharides (LPS) extracted from the Gram-negative bacterial cell walls are complex mixtures with significant structural heterogeneity. ^[1-3]^ The three main components of an S-type LPS (smooth LPS) are lipid A, core oligosaccharide, and O-antigen. R-type LPS (rough LPS) is missing the terminal O-antigen and therefore consists of only lipid A and core oligosaccharide (LOS). The complexity of the LPS mixtures arises from variations in the number, type, and positions of fatty acids and sugars in their chemical structures. Moreover, variations in the number and position of additional functional moieties, such as phosphate, acetyl, and phosphoryl ethanolamine groups, add more complexity to the LPS structural heterogeneity. Thus, fractionation and separation of individual components of the LPS mixtures are extremely difficult using the previously tested chromatographic methods. ^[1 and references therein]^ For this reason, only a few studies to date have chosen to use liquid chromatography-tandem MS (LC-MS/MS) for the analysis of intact LPS or LOS complex mixtures ^[4-5]^

The existence of most LPS and LOS ions as mixtures of isomers and/or isobars complicates their structural elucidation by tandem MS. ^[3-9]^ The co-isolation of isomeric and isobaric ions for fragmentation results in chimeric spectra that are difficult to interpret.^[10,11]^ For this reason, ion mobility MS is an excellent choice for separating the expected LPS and LOS isomeric and isobaric ions prior to tandem MS. Field Asymmetric Ion Mobility Spectrometry (FAIMS) offers gas-phase fractionation of ions based on their charge states and mobility differences in the strong and weak electric fields. ^[12-14]^ The FAIMS fractionation mechanism is ideal for LPS/ LOS analysis as it can eliminate singly charged chemical background ions. ^[15]^ These singly charged ions suppress the expected LPS/LOS multi-charge ions intensities hindering their characterization by tandem MS. Previously, capillary electrophoresis coupled to FAIMS has been reported for the analysis of the O-deacylated *Haemophilus influenzae* LOS prepared by treating the intact LOS with hydrazine. ^[15]^ Thus, this is the first report on the use of FAIMS to analyze an intact or untreated LOS complex mixture with the ability to resolve isobaric LOS ions and their characterization by tandem MS.

LPS structural elucidation by MS was traditionally facilitated by chemical extraction of either the lipid A or the oligosaccharide and polysaccharides components, followed by additional chemical degradation and/or modification. ^[16-25]^ However, chemical degradation can lead to the loss of crucial structural information and thus interferes with attempts to understand the original LPS or LOS intact structures present *in vivo*. While a few structural studies have been done on intact LPS or LOS mixtures without any chemical treatment, ^[4-9]^ no in-depth or comprehensive profiling of the intact LPS or LOS mixtures has been reported.

Kendrick mass defect (KMD) plots are valuable tools for classifying and sorting structurally related ions in complex mixtures. ^[26-28]^ Traditional KMD plots use the methylene group base unit [CH_2_] to classify ions, especially in petroleomics and lipidomics. ^[29,30]^ KMD plots represent a relationship between the Kendrick mass defect (KMD) and the exact Kendrick mass (KM). The KM (Eq.1) and KMD (Eq.2) are calculated as follows:

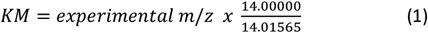

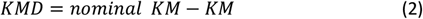

All ions with the same chemical composition but different number of CH_2_ will have the same Kendrick mass defect and will align on the same horizontal line in the KMD plot, thus creating a KMD ion family. On the other hand, ions aligning on the same vertical and/or diagonal line usually differ in the number of double bond equivalents (DBE or degree of unsaturation) or the number of heteroatoms; i.e., the ion with a higher DBE or a number of heteroatoms will have a higher Kendrick mass defect. ^[26-28]^

Based on the diagnostic structural features of the studied class of compounds, other custom KMD base units instead of the traditional CH_2_ can be employed to sort structurally related ion families. ^[27,28]^ In this case, the modified exact Kendrick mass (Eq.3) is calculated as follows:

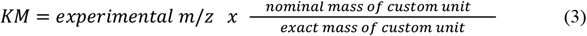

Since structurally related ions always align on the same horizontal and/or vertical lines of the KMD plot (KMD ion family), KMD plots allow better visualization of the structural relationship between different ions and the in-depth profiling of complex mixtures. During data analysis, KMD plots, therefore, reduce the number of tandem mass spectra needed to characterize each KMD ion family. By identifying the structural relationships between ion families across a KMD plot, confirming a specific ion’s structure by tandem MS can be used to propose the chemical composition of other structurally related ions. ^[27,28]^

This work showcases the benefits of FAIMS rapid gas phase fractionation and KMD plots in the characterization of *E. coli* J5 (Rc mutant) LOS, a well-known TLR4 agonist used previously as a vaccine adjuvant. ^[31]^ Analysis was accomplished using direct infusion FAIMS-electrospray ionization (ESI)-tandem MS followed by KMD plots analysis using a custom [Na-H] base unit. This combined approach allowed the in-depth characterization of 179 LOS elemental compositions out of 321 present in the mixture and the identification of novel structural features of the J5 *E. coli* (Rc mutant) LOS.

## 2. Materials and Methods

### 2.1. Direct infusion FAIMS-ESI-tandem MS

Negative ion mode ESI-MS was performed on an Orbitrap Fusion ETD Tribrid Mass Spectrometer with a FAIMS pro interface (Thermo Scientific, Waltham, MA, USA). Rough LOS of *E. coli* J5 Rc mutant (Sigma Aldrich, St. Louis, MO, USA) was dissolved in MS-grade water: isopropanol: triethyl amine: acetic acid (80:20:0.2:0.2 v/v/v/v) at a concentration of 1 μg/μL and directly infused into the FAIMS source at a flow rate of 3 μL/min. The spray voltage was set to 2.5 kV, capillary temperature to 275 °C, and sheath and auxiliary gas to 2 a.u. The FAIMS separation was accomplished with the following settings: inner electrode temperature = 100 °C, outer electrode temperature = 100 °C, FAIMS N_2_ carrier gas flow = 1.2 L/min, asymmetric waveform with dispersion voltages (DV) = +5000 V. Compensation voltages (CV) were scanned from 0 to 100 V with a step size of one volt at 120 K resolution, 150 – 2000 *m/z* range, 60% RF lens, 100% normalized automatic gain control (AGC) target, and 100 ms Maximum injection time (IT). The mass spectrum was also acquired with the FAIMS off mode for comparison purposes. Tandem MS was performed at selected CV voltages (50 and 61 V) that separate isobaric ions of interest (*m/z* 1085.17 and *m/z* 1085.20). Normalized high-energy collision-induced dissociation (HCD) in the 30-40% range was used for tandem MS. Pseudo-MS^3^ was accomplished with source-induced dissociation (SID) at a potential difference of 100 V. The source-induced dissociation fragmented LOS precursor ions into lipid A (Y-type fragments) and core oligosaccharide fragments (B-type fragments). The resulting B-type fragments of interest were further selected for HCD MS/MS (Pseudo MS^3^)

### 2.2. Data analysis

The Mass spectra at different FAIMS compensation voltages were viewed and deconvoluted using the *Xtract* tool in Freestyle software 1.8 SP1 (Thermo Scientific). The deconvolution parameters were: 2-4 charge range, 1% threshold relative abundance, protein isotope table, and a minimum number of detected charges of 1. Any masses below the molecular weight of 2237.34 Da corresponding to Kdo2-Lipid A (KLA) were removed. The deconvoluted mass list from all CVs combined (average of 100 scans) was uploaded to the Constellation online software (https://constellation.chemie.hu-berlin.de/). ^[32]^ A custom KMD base unit of [Na-H] corresponding to the exact mass of 21.981945 Da was used to create the KMD plot. One of the LOS ions annotated as ***24*** in the KMD plot (**Figure S6**) was the key to predict the chemical composition of all the other LOS ions.

The chemical composition of ion ***24*** was determined by a detailed tandem MS characterization (MS^2^ and Pseudo MS^3^). Tandem MS fragmentation was investigated manually with the assistance of the ChemDraw software v. 21.0.0.28 (Perkin Elmer, Waltham, MA, USA) and the GlycoWork bench software 1.1.3480. ^[33]^ Structural assignments were based on previously reported J5 Ecoli LOS proposed structures and/or biosynthesis. ^[34-36]^ Similar dots arrangement and/or geometric pattern of different LOS ion families across the KMD plot allowed the determination of chemical composition of extra 73 LOS ions with reference to the ion ***24*** chemical formula elucidated by tandem MS. **Figure S6** shows the chemical formulas of ions 24, 25, and 19 as a representative example of chemical formula assignments. The alignment of ion 25 on the same vertical line with ion ***24*** indicates that ion ***25*** has an extra phosphate group compared to ion ***24***. Thus, ion ***25*** has the same chemical formula as ion ***24*** plus HPO_3_ (**Figure S6**). Similarly, difference of 198.16 Da between ions ***19*** and **24** corresponds to C12:0 acyl chain plus one oxygen (**Figure S6**). Therefore, ion 19 has the same chemical formula as ion ***24*** minus C_12_H_22_O_2_. All chemical composition assignments for the identified 73 LOS ions were within +/-5 ppm (**Table S2**).

## 3. Results and Discussion

### 3.1. LOS characterization by Direct Infusion ESI-FAIMS-tandem MS

In this study, we have investigated the benefits of FAIMS for analysis of the complex mixture of *E. coli* J5 (Rc mutant) LOS composed of lipid A and core oligosaccharide (R-type LPS). When FAIMS compensation voltages were scanned from 0-100 volts, gas fractionation of LOS ions was achieved, allowing the identification of a higher number of multi-charged LOS ions (**Figure 1**). ^[37,38]^ In contrast, when the FAIMS voltages were not applied (FAIMS-off mode), many of the LOS multi-charged ions were suppressed in favor of singly charged background ions and phospholipid contaminants (**Figure S1**). The number of LOS species (> 2237.34 Da) that resulted from compensation voltage fractionation increased by approximately 1.7-fold compared to when FAIMS voltages were off (191 vs. 321 deconvoluted masses). Moreover, the use of specific FAIMS compensation voltages, such as CV=57, allowed the elimination of singly charged ions and enhancement of the LOS ion signal-to-noise (S/N) ratio (**Figure S1**). A representative example is the S/N ratio enhancement of the LOS ion at *m/z* 940 by approximately 2.4-fold at CV=57 compared to its S/N ratio when the FAIMS voltages were off. (**Figure S1**).

**Figure 1:**
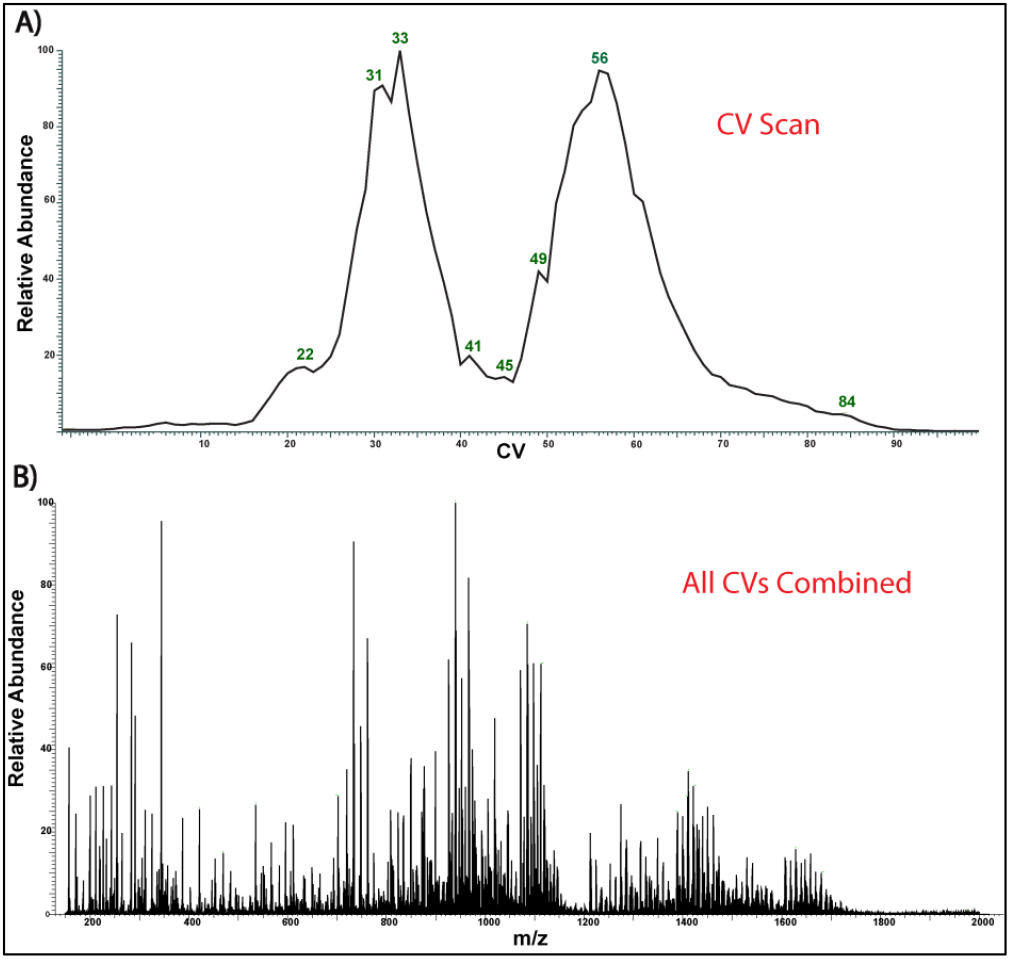
FAIMS-ESI-MS of the J5 *E. coli*. A) FAIMS Compensation voltage scan (0-100 V). B) Corresponding spectrum of all CVs combined (average of 100 scans)

The presence of isobars and overlapping isotopic distributions often hinder the accurate structural characterization of ions of interest by tandem MS. When FAIMS compensation voltages (CV) were scanned from 0 to 100 V; it was noted that several overlapping LOS isobaric ions were separated at specific compensation voltages (**Figure S2**). For example, the most intense LOS ion at *m/z* 1085 (*z* = -3) detected when FAIMS voltages were off showed two overlapping isobars, a major ion at *m/z* 1085.17 and a minor ion at *m/z* 1085.20. This overlap was still notable at different compensation voltages, such as CV=60, when FAIMS voltage was on (**Figure S2A**); however, at CV=61, only the minor ion at *m/z* 1085.20 was detected (**Figure S2C**). Furthermore, at CV=50, the major isobar at *m/z* 1085.17 was separated from any traces of the minor ion (**Figure S2B**). Therefore, carefully manipulating the FAIMS CV allowed for efficient separation of the most intense LOS isobaric ions at *m/z* 1085.17 and 1085.20 for a follow-up tandem MS, avoiding the generation of chimeric mass spectra.

Previous tandem MS structural characterization of *E. coli* J5 (Rc mutant) LOS by Oyler *et al*. showed that the chemical formula of the triply charged minor ion at *m/z* 1085.20 was [C_145_H_259_N_3_O_70_P_3_]^3-^.^[7]^ This LOS ion was proposed to be comprised of the canonical hexa-acylated *E. coli* lipid A ^[24,25]^ and a core oligosaccharide (OS) consisting of seven monosaccharides (two 3-deoxy-D-manno-oct-2-ulosonic acid: Kdo, three heptose: Hep, one hexose: Hex and one glucosamine: GlcN) and modified with one phosphate (P) and one acetyl group (Ac): Kdo2. Hep3. Hex. GlcN. P. Ac. ^[7]^ In the current study, tandem MS analysis of this minor LOS ion at *m/z* 1085.20 (*z* = -3) (**Figure 2A**) yielded two main product ions: *m/z* 1796.20 (*z* = -1) and *m/z* 1240.33 (*z* = - 1), corresponding to lipid A (Y-type product ion) and oligosaccharide [OS-H_2_O-Kdo] (B-type product ion), respectively, according to the Domon and Costello nomenclature. ^[39]^ The B-type product ion at *m/z* 1240.33 was detected mainly as a singly charged ion due to the presence of a single phosphate group in its chemical composition, confirming the previous findings by Oyler *et al*. (**Figure 2A**). ^[7]^

**Figure 2:**
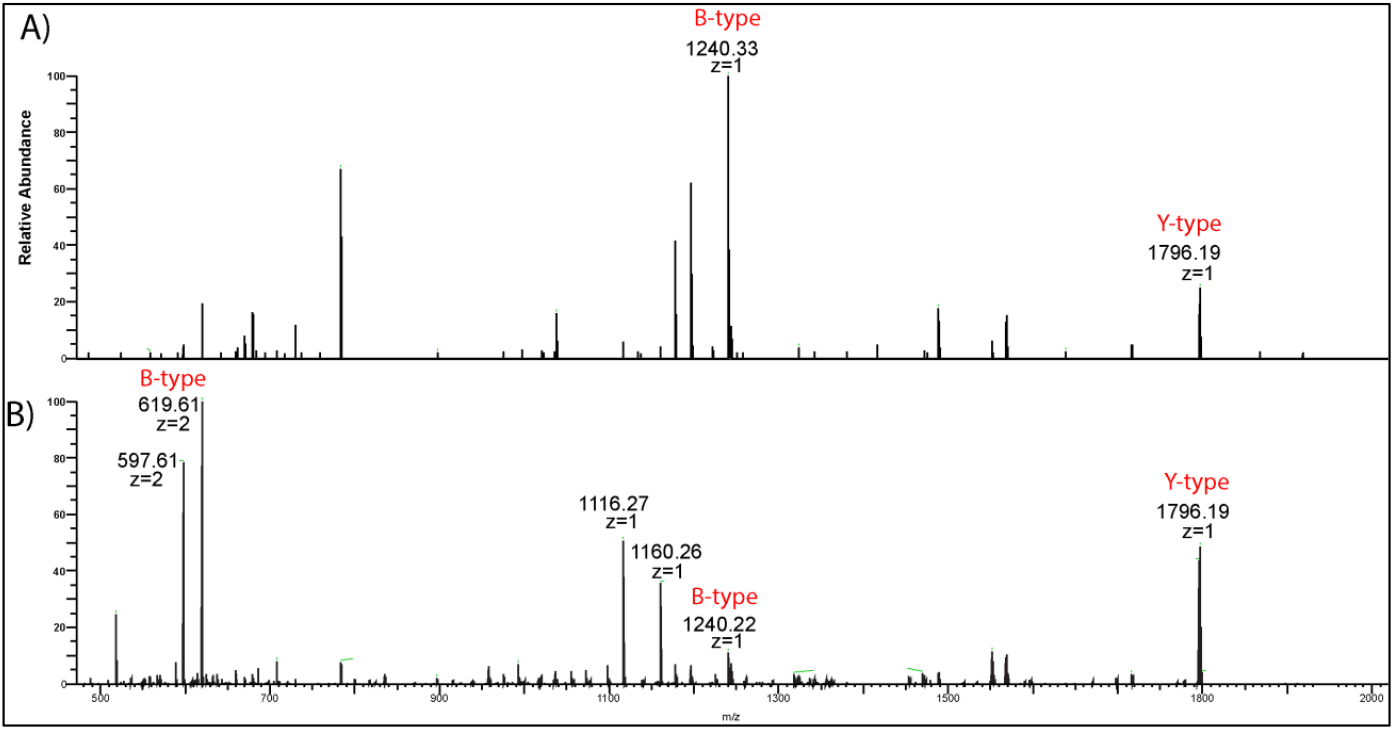
**A)** High-energy collision-induced dissociation HCD MS/MS of **a)** *m/z* 1085.20 isolated at CV=61. **B)** *m/z* 1085.17 isolated at CV= 50. Both spectra showed the formation of the same Y-type ion (*m/z* 1796), but the respective B-type ions differed in their exact masses

The tandem MS analysis of the major LOS ion at *m/z* 1085.17 (**Figure 2B**) also produced the same lipid A Y-type product ion at *m/z* 1796.20 (*z* = -1). However, the corresponding OS B-type product ion showed a different exact mass (*m/z* 1240.23) and lower intensity compared to the one detected for the minor ion (*m/z* 1240.33; **Figure 2A**). This latter B-ion at *m/z* 1240.23 showed the loss of a phosphate group (−80 Da) to give the ion at *m/z* 1160.26. Further loss of CO_2_ (−44 Da) from the Kdo sugar resulted in the ion at *m/z* 1116.27. Importantly, the same B-ion at *m/z* 1240.23 (*z* = -1) was also detected as a highly intense doubly charged ion at *m/z* 619.61 (*z* = -2) that showed a loss of CO_2_ to form the ion at *m/z* 597.61 (*z* = -2) (**Scheme 1**). Based on this fragmentation pattern and the presence of the highly intense doubly charged B-ion, we concluded that the major LOS (*m/z* 1085.17) ion should have at least two phosphate groups in its core oligosaccharide.

From this analysis, tandem MS alone could only suggest the number of phosphate groups in the major LOS ion but not its monosaccharide composition or sequence. For this reason, MS^3^ was needed to elucidate the possible monosaccharide composition and sequence of the oligosaccharide. ^[40]^ Source-induced fragmentation (SID) of 100 V was utilized to create the intense doubly charged B-type ions that were further selected for tandem MS (Pseudo MS^3^). Tandem MS of the B-type ion at *m/z* 597.61 that dominated the SID mass spectrum (**Figure S3**) represented the loss of a phosphate group resulting in the major singly charged ion at *m/z* 1116.27. This loss indicates the instability of the selected doubly charged ion, hindering the identification of the monosaccharide from which this phosphate group has been lost **(Figure 3)**. However, previously reported *E. coli* J5 biosynthesis suggests that this phosphate group is on C-4 of the second heptose moiety denoted as Hep II (**Scheme 1**). ^[34-36]^

**Figure 3:**
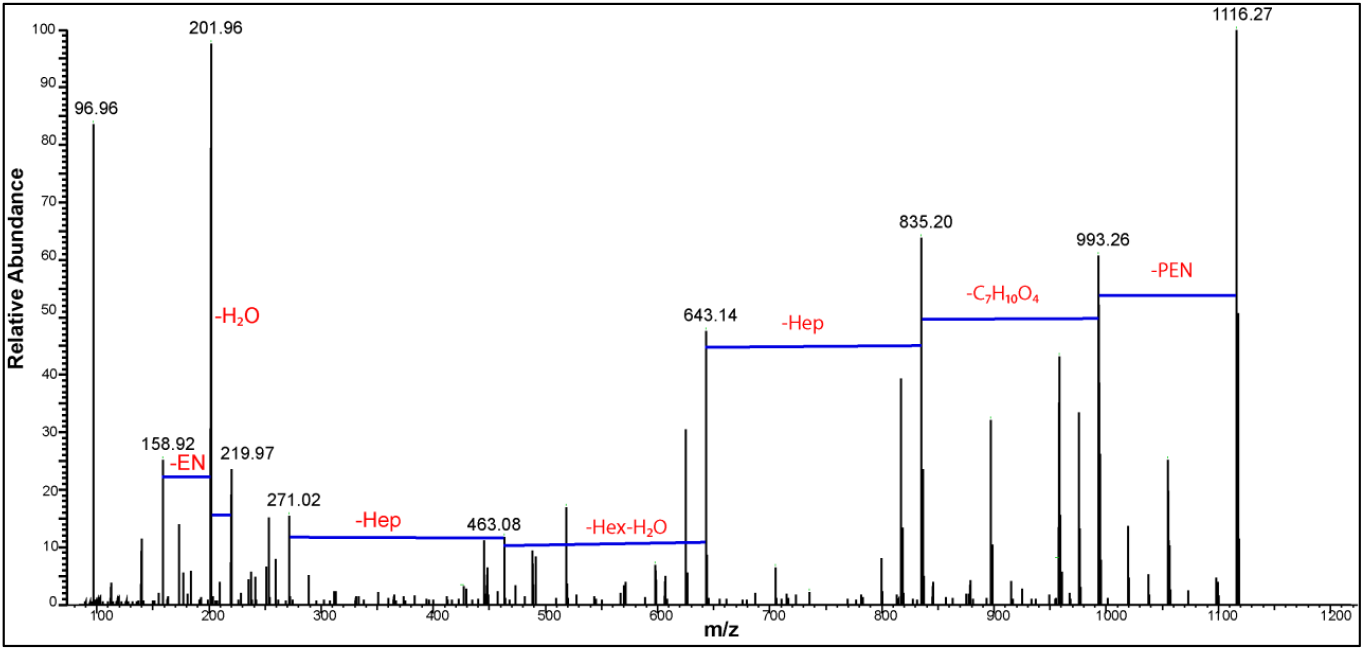
HCD MS/MS (Pseudo MS^3^) of *m/z* 597.61 created by source induced dissociation of 100 V

Also, the double-charged ion at *m/z* 597.61 decomposed to create two singly charged ions at *m/z* 201.97 and *m/z* 993.26. It should be noted that ions in the low mass region at *m/z* 219.97, 201.96 and 158.92 confirm the presence of pyrophosphoryl ethanolamine group **(Figure 3)**. Also, another pathway to create *m/z* 993.26 is the loss of phosphoethanol amine group (−123 Da) from the ion at *m/z* 1116.27. Thus, *m/z* 993.26 is represented with two isomeric forms **(I and II)** in **Scheme 1**. In addition, note that the PPEN group was placed on the C-4 of the first heptose sugar (Hep I) to harmonize with the known *E. coli* biosynthetic pathways. ^[34-36]^

The singly charged key ion at *m/z* 993.26 showed a series of sequential losses highlighting its monosaccharide components. The first was the loss of 158.05 Da corresponding to C_7_H_10_O_4_, and equivalent to the chemical composition of [Kdo-H_2_O-CO_2_]. This loss formed the ion at *m/z* 835.21, which lost a heptose sugar (−192 Da) to produce *m/z* 643.14. The last ion at *m/z* 643.14 experienced the combined loss of a hexose and water molecule (−180 Da) to create *m/z* 463.08, which in turn lost a heptose moiety (−192 Da) to produce *m/z* 271.02 corresponding to a phosphorylated heptose. Other ions confirming the proposed structure of *m/z* 993.26 are annotated in the pseudo MS^3^ of *m/z* 597.61 (*Z* = -2) (**Figure S4**) and listed in **Table S1**.

The combined fragmentation patterns from the MS^2^ of *m/z* 1085.17 and the pseudo MS^3^ of the doubly charged B-type ion at *m/z* 597.61 showed that OS was composed of six sugars modified with one phosphate and one pyrophosphoryl ethanolamine: Kdo2. Hep3. Hex. P. PPEN. **Figure S5** shows the full structure of the ion at *m/z* 1085.17 and how the Y-type (lipid A) and B-type ion fragments (OS) were created.

With such complex, non-polymeric compounds, manual tandem MS data analysis was time-consuming and not a straightforward task. First, neutral and/or charged losses (**Scheme 1**) at different tandem MS stages (**Figures 2 and 3**) had to be carefully inspected to deduce product ions genealogy. This complexity was clearly demonstrated by the presence of two possible fragmentation pathways to form the key ion *m/z* 993.26 **(Scheme 1)**. Also, one of the apparent complexities is the presence of two main modifications (i.e., phosphate and pyrophosphryl ethanolamine) and six monosaccharides in the B-ion chemical structure (**Scheme 1**). Thus, several possible structures (6^2^ = 36 possibilities) must be investigated to find the best possible structure. These challenges were the driving force for explaining the detailed tandem MS fragmentation mechanisms (**Scheme 1**) (*vide supra*) to serve as a simplified tutorial for the LPS tandem MS characterization.

**Scheme 1:**
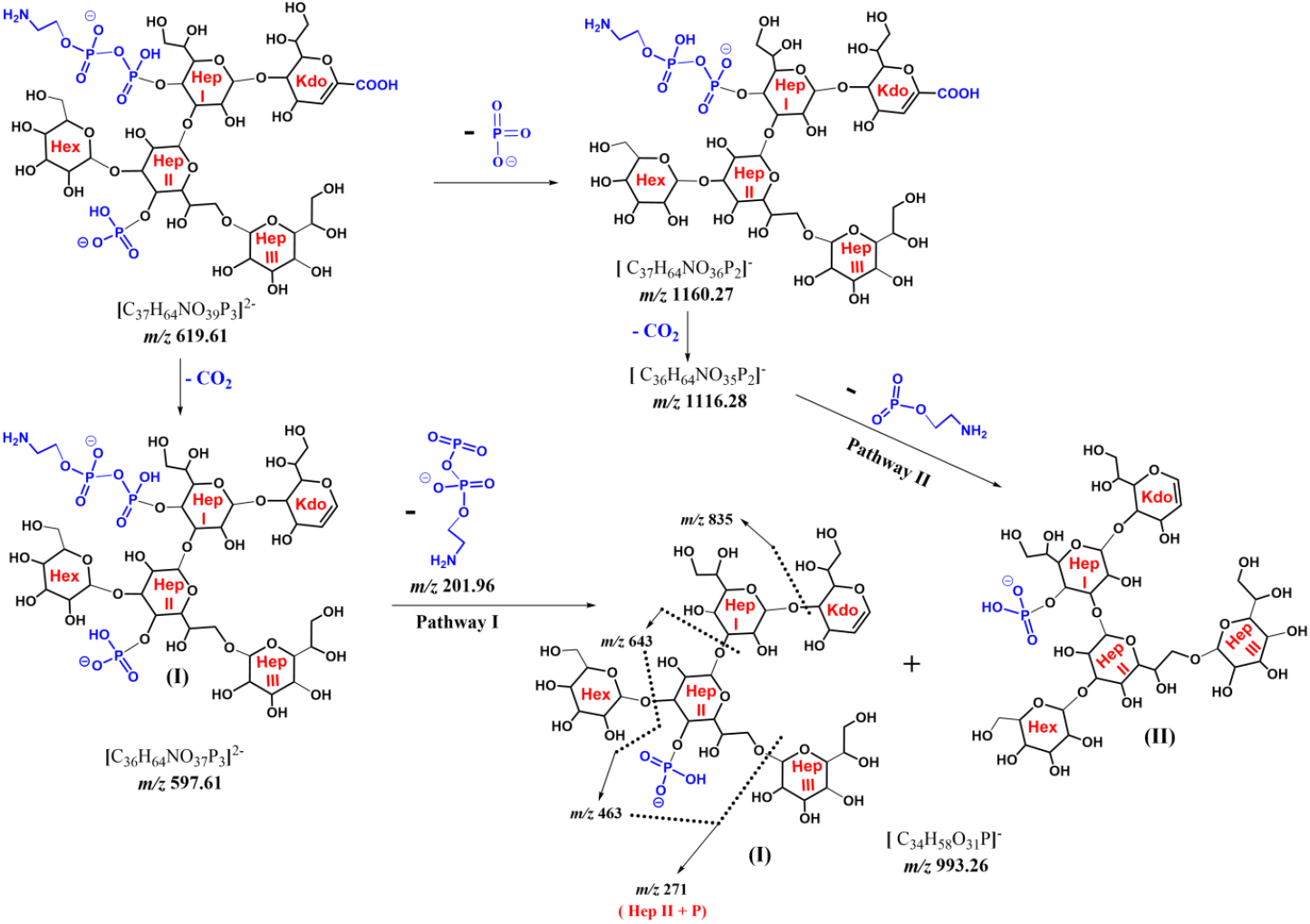
MS^2^ and Pseudo MS^3^ fragmentation mechanism of B-ions originating from *m/z* 1085.17.

### 3.2. In-depth LOS characterization by Kendrick mass defect (KMD) plots

The previous section demonstrated that analysis of LOS structures by tandem MS data is challenging and complicated by the lack of software for automated data analysis. Additionally, there are no libraries of possible structural variations between the LPS ions that would facilitate tandem MS analysis of LPS or LOS.

Therefore, Kendrick mass defect plots were used to streamline the analysis of structural variations across the different LOS species. The LOS ions were detected mostly as deprotonated ions with the formula (M - zH)^-z^ along with some minor sodiated clusters with the formula [M – zH + n (Na - H)]^-z^ (n =1, 2 and 3). Hence, a highly organized KMD plot was created with [Na-H] base unit of 21.981945 Da. ^[27,28]^ In this plot, ions with different numbers of sodium ions were arranged on horizontal lines, while ions with varying numbers of phosphate groups were arranged on diagonal lines (**Figure 4**). The alignment of sodiated clusters on the same horizontal line allows the visual differentiation between the major unsodiated ions and their sodiated analogs. Thus, it facilitates identifying structural relationships between the different LOS ion families across the KMD plot.

**Figure 4:**
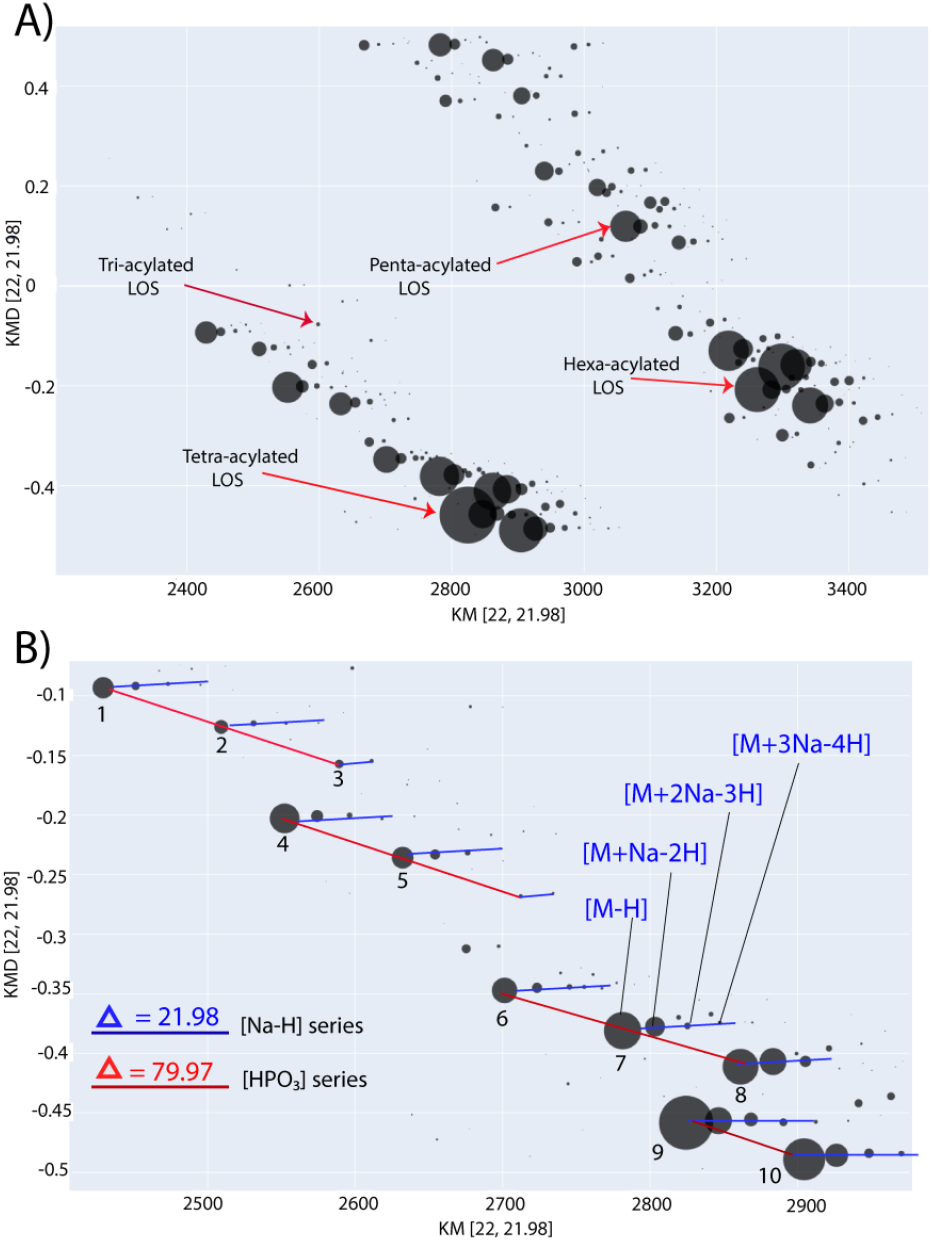
KMD plot of the *E. coli* J5 (Rc mutant LOS) using [Na-H] base unit. **a)** Simplified KMD plot showing LOS families with different acylation degrees in the molecular weight range of 2200-3600 Da and their relative ion intensities. **b)** KMD plot region showing horizontal ion series (blue lines) differ by 21.98 Da [Na-H] while vertical ion series (red lines) differ by 79.97 [HPO3]. Ion clusters **(1-5)** have similar KMD arrangement/geometric pattern as ions cluster **(6-10)** demonstrating a constant structural difference of 272.03 Da corresponding to the combined incorporation of one phosphate and one heptose.

Moreover, several ion families had similar geometric patterns across the KMD plot; e.g., the ion families annotated as ***1-5*** (**Figure 4B**, in blue) and ***6-10*** (**Figure 4B**, in red). Careful visual inspection of these similar geometric patterns allowed for the determination of interesting structural differences and/or relationships between the different LOS ion families. For example, the mass difference between individual ions of the two families **(*1-5*** and ***6-10*)** was 272.03 Da; thus, the difference between ions ***6*** and ***1*** is the same as the difference between ions ***2*** and ***7***, and so forth. This 272.03 Da difference can be explained by adding a single phosphorylated heptose or adding one heptose and one phosphate to different positions of the core oligosaccharide. This may indicate that this ion cluster ***(1-5)*** represents a set of structures that form a defined biosynthetic modification pathway that created a new ion family ***(6-10)***. An addition of a phosphorylated heptose is more likely, in this case previously described in *E. coli*. ^[41,42]^ Briefly, heptosyl transferases (Waac, Waaf and WaaQ) are responsible for the addition of heptose sugars, while LOS kinase (WaaP) and LOS phosphatase (WaaY) add phosphate groups to heptoses. ^[41,42]^

Besides the core oligosaccharide structural differences, various degrees of acylation in the lipid A moiety were detected from KMD structural network: hexa-acylated, penta-acylated, tetra-acylated, and minor tri-acylated LOS ion clusters. ^[4]^ As an example of acylation heterogeneity, **Figure S6** shows the difference of 198.16 Da corresponding to C12:0 (182 Da) plus oxygen (16 Da) between ions cluster ***(16-20)*** and ***(21-25)*** representing penta-acylated and hexa-acylated LOS species, respectively.

Remarkably, the detailed structural elucidation of a single ion ***24*** (*m/z* 1085.17) by tandem MS allowed the prediction of chemical compositions of other 72 LOS ions (**Table S2**) with the help of the KMD structural network as explained previously in detail in the data analysis method (**Section 2.2**). This clearly shows the usefulness of KMD plots in reducing the number of challenging tandem mass spectra that need to be interpreted manually. It also assists in proposing possible chemical compositions of low abundant ions (***35-40*, Figure S7**) that are challenging to characterize by tandem MS. The identified structural differences could be in turn, used to characterize mechanisms of LPS/LOS biosynthesis and modification. The detailed knowledge about LPS biosynthesis, modifications, and structural variations will definitely allow an in-depth understanding of the function of the Gram-negative outer membrane. ^[43]^

Interesting structural differences (**Figure S8**) were also detected, such as 81.10 Da indicating the replacement of a phosphate group with a glucosamine sugar between ion clusters ***(21-25)*** and ***(31-35)***. Differences in the number of phosphoryl ethanolamine and/or ethanolamine groups were also identified, as a few minor ions contained up to three ethanolamine groups. Other minor differences were acetylation (+42.01 Da) and missing C_2_H_4_ units of the lipid A (−28.03 Da), indicating the presence of a shorter acyl chain. Moreover, a difference of 42.03 Da were detected between two minor ions, which is likely to be an addition of C_3_H_6_ (trimethylation), indicating either a longer lipid A acyl chain and/or methylation of the core oligosaccharide. All the structural differences deciphered from the KMD structural network are listed in **Table 1**. The previously described examples demonstrate how the created KMD plot can provide meaningful insights into the structural relationships and/or organization between the different LOS species.

**Table 1:** Structural differences deciphered form the KMD plot analysis of the J5 E. coli LOS (Rc mutant). Abbreviations used are Heptose: Hep, Glucosamine: GlcN, phosphate: P, ethanolamine: EN, Oxygen: O, 3-hydroxy-tetradecanoic acid: C14:0(3-OH), dodecanoic acid: C12:0, tetradecanoic acid: C14:0

| Structural Difference (Da) | Corresponding structure |
| --- | --- |
| 41.91 | PEN+P-GlcN |
| 42.01 | Ac |
| 42.04 | C<sub>3</sub>H<sub>6</sub> |
| 43.03 | EN |
| 44.02 | CH<sub>2</sub>CH<sub>2</sub>O |
| 45.84 | Hep + P - [C14:0(3-OH)] |
| 73.87 | Hep + P - [C12:0 + O] |
| 79.96 | P |
| 81.10 | GlcN - P |
| 121.87 | PEN+ 2P-GlcN |
| 123.00 | PEN |
| 157.12 | [C14:0]+ C<sub>2</sub>H<sub>4</sub> + GlcN- P |
| 164.36 | [C14:0] + [C14:0(3-OH)] - Hep- P |
| 198.16 | [C12:0 + O] |
| 238.23 | [C14:0] + C<sub>2</sub>H<sub>4</sub> |
| 272.03 | Hep + P |
| 307.30 | [C14:0(3-OH)] + GlcN - P |
| 353.14 | Hep + GlcN |
| 390.55 | [C14:0] + [2 x C14:0(3-OH)] - Hep- P |
| 436.40 | [C14:0] + [C14:0(3-OH)] |
| 464.42 | [C14:0] + [C14:0(3-OH)] + C<sub>2</sub>H<sub>4</sub> |
| 510.26 | [C14:0]+ C<sub>2</sub>H<sub>4</sub> + Hep+ P |
| 517.50 | [C14:0] + [C14:0(3-OH)] + GlcN - P |
| 662.58 | [C14:0] + [2 x C14:0(3-OH)] |
| 708.42 | [C14:0] + [C14:0(3-OH)]+ Hep + P |
| 743.68 | [C14:0] + [2x C14:0(3-OH)] + GlcN - P |
| 789.52 | [C14:0] + [C14:0(3-OH)] + Hep + GlcN |

Lastly, it should be mentioned that the identified 73 LOS ions (**Table S2**) were comprised of 23 hexa-acylated LOS, 15 penta-acylated LOS, 27 tetra-acylated LOS, and 8 tri-acylated LOS. In addition, the corresponding sodiated clusters of each of these 73 ions (ions arranged on the same horizontal line with the main unsodiated ion, Figure 4B) can also be identified, bringing up the total identified chemical compositions to 179 LOS species. Moreover, the relative signal intensity between the most intense ion from each LOS family (annotated in **Figure 4A**) was approximately 12:8:15:1 for hexa-acylated, penta-acylated, tetra-acylated, and tri-acylated LOS, respectively. This analysis allowed the various structures from likely agonistic to antagonistic relative to the Toll-like receptor 4 (TLR4)/myeloid differentiation factor 2 (MD-2) complex to be estimated. The comprehensive information provided by the KMD plot detailing the LOS acylation degree and/or heterogeneity may lead to an increased understanding of the structure-immunoactivity relationship of LOS and innate immune responses to TLR4 agonists.

## 4. Conclusion

We presented here the valuable benefits of using FAIMS rapid gas-phase fractionation and KMD plots for intact LOS characterization. The combined use of ESI-FAIMS-MS^/^MS and KMD plots allowed the in-depth characterization of the *E. coli* J5 (Rc-mutant) LOS complex mixture. The FAIMS isobaric ion separation facilitated LOS detailed structural elucidation by tandem mass spectrometry. On the other hand, the custom KMD plot using [Na-H] offered a great tool for visualizing the structural relationships between the different LOS species. The ions organized arrangements across the KMD plot, and the detailed tandem MS characterization of one ion allowed the determination of 73 unique LOS ion compositions in addition to their sodiated analogs. This approach allowed the development of a list of interesting structural differences that may highlight a biosynthetic step and/or explain the biological activity behind the current and future *E. coli* LOS-based glycoconjugate vaccine adjuvants.

The ESI-FAIMS-MS/MS and KMD plot approach and the data presented here can serve as a foundation for creating a library of structural differences for LPS from various bacterial strains. This developed data and methods used herein will enhance development of software utilizing the structural differences list revealed by KMD plots for automated MS data analysis. Future work will focus on analyzing other rough or smooth types of LPS from different bacterial strains. The main target is to develop simple methods and/or structural libraries to facilitate more efficient characterization of LPS complex mixtures.

## Supporting information

LPS_FAIMS_KMD_Supplementary figures and tables

## Funding and Acknowledgement

DRG and RKE thanks the National Institutes of Health for grant R01 AI123820-01. DRG thanks Genome BC through the Sector Innovation Program (SIP7) for support,

## Notes

### Competing Interest Statement

The authors have declared no competing interest.

### Summary of Updates

Scheme 1 revised and more supplemental information added to support the proposed chemical structures

