## Supplementary material for "Structural Elucidation of Intact Rough-Type Lipopolysaccharides using Field Asymmetric Ion Mobility Spectrometry and Kendrick Mass Defect Plots": LPS_FAIMS_KMD_Supplementary figures and tables

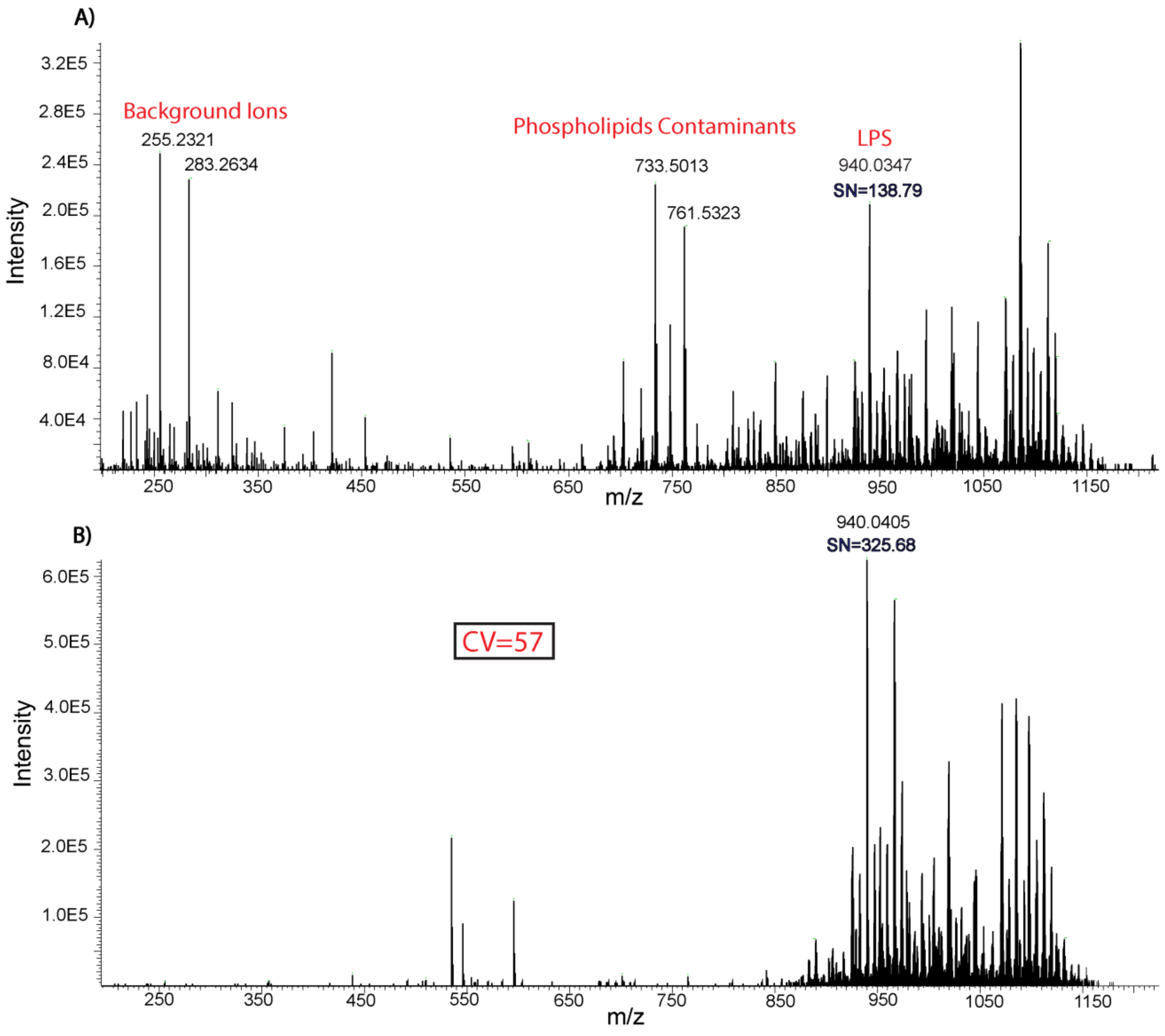


**Figure S1:** **A)** ESI-MS of the J5 *E. coli* LPS (FAIMS off). **B)** Corresponding mass spectrum at CV=57 showing the S/N ratio enhancement of *m/z* 940.04 ratio by 2.4 fold compared to when the FAIMS voltages were off

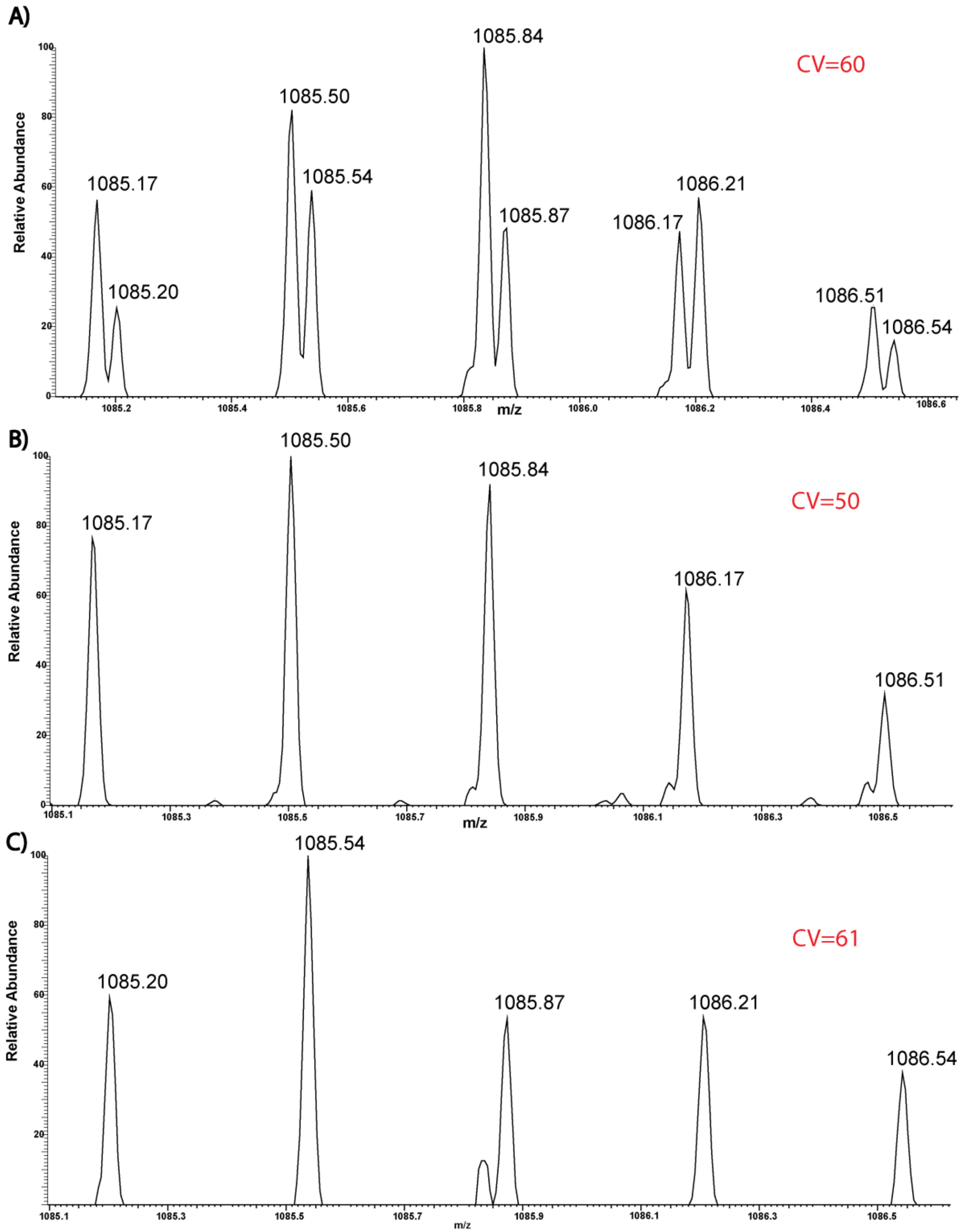


**Figure S2:** Separation of LPS isobaric ion at *m/z* 1085. **a)** Overlapping isobars at *m/z* 1085.17 and 1085.20 at CV=60 V. **b)** Separated isobar at *m/z* 1085.17 at CV=50 V. **c)** Separated isobar at *m/z* 1085.20 at CV= 61V.

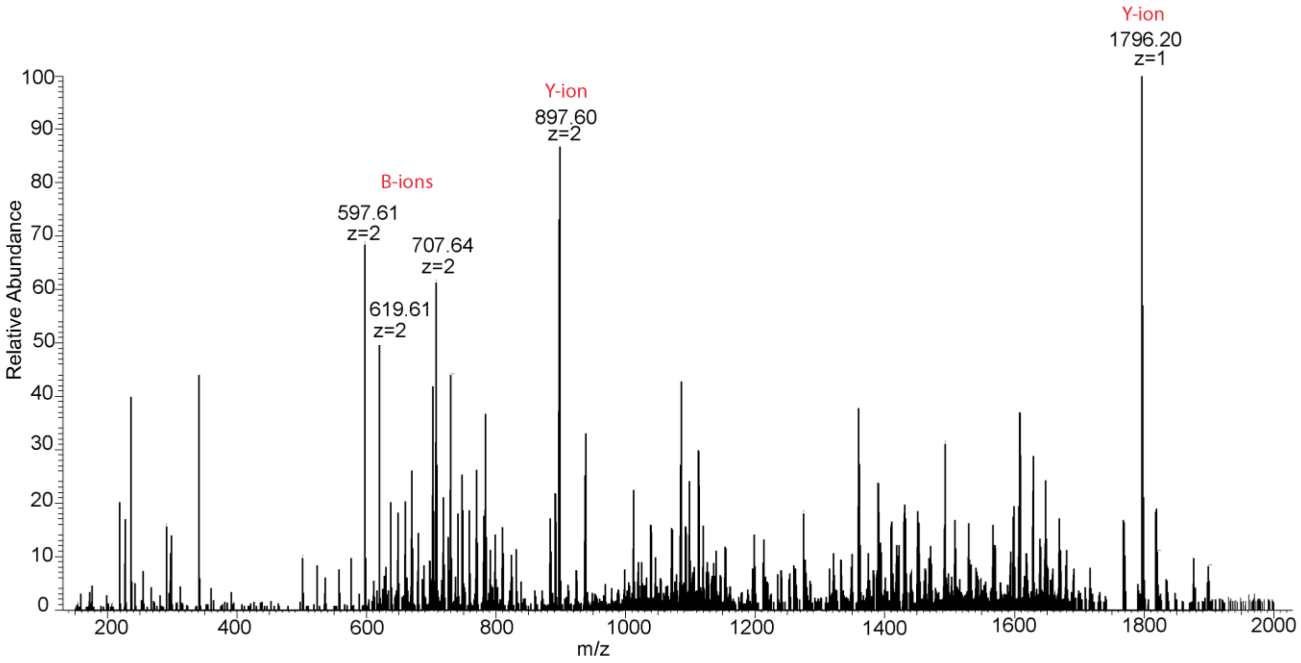


**Figure S3**: Source induced dissociation (SID) of *E. coli* J5 LPS. Fragmentation of the LOS ions in the source were accomplished by applying an energy of 100 V producing *m/z* 597.61 corresponding to the major oligosaccharide fragment of *m/z* 1085.17.

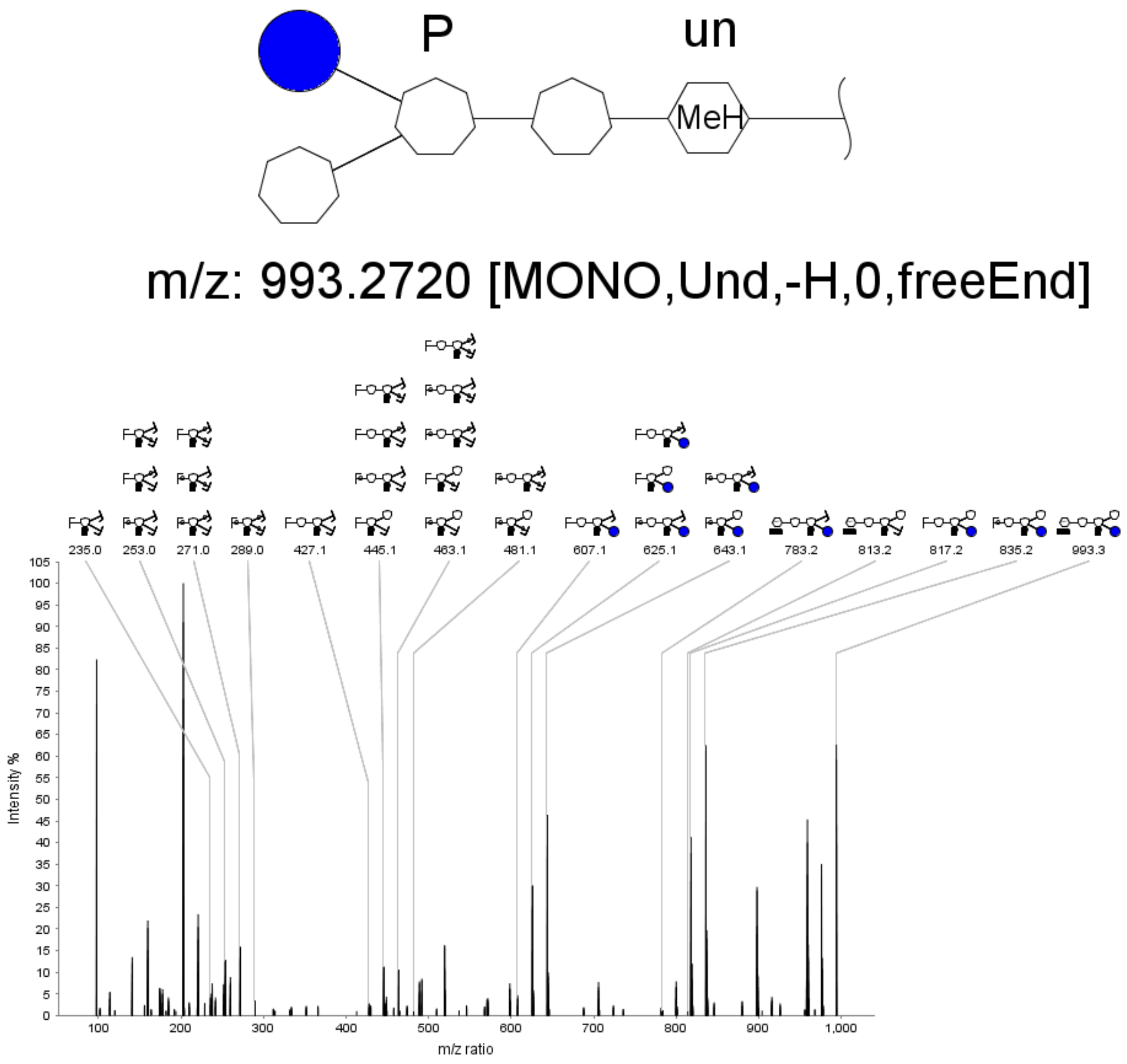


**Figure S4**: Annotated pseudo $MS^3$ of *m/z* 597.61 ($Z$ = -2) generated using the GlycoWork bench software showing product ions originating from the singly charged key ion at *m/z* 993.26. The [Kdo-$H_2O$-$CO_2$] or $C_7H_{10}O_4$ residue in *m/z* 993.26 is represented in the software by an unsaturated methyl hexose residue [$C_7H_{12}O_5$-$H_2O$ = $C_7H_{10}O_4$]

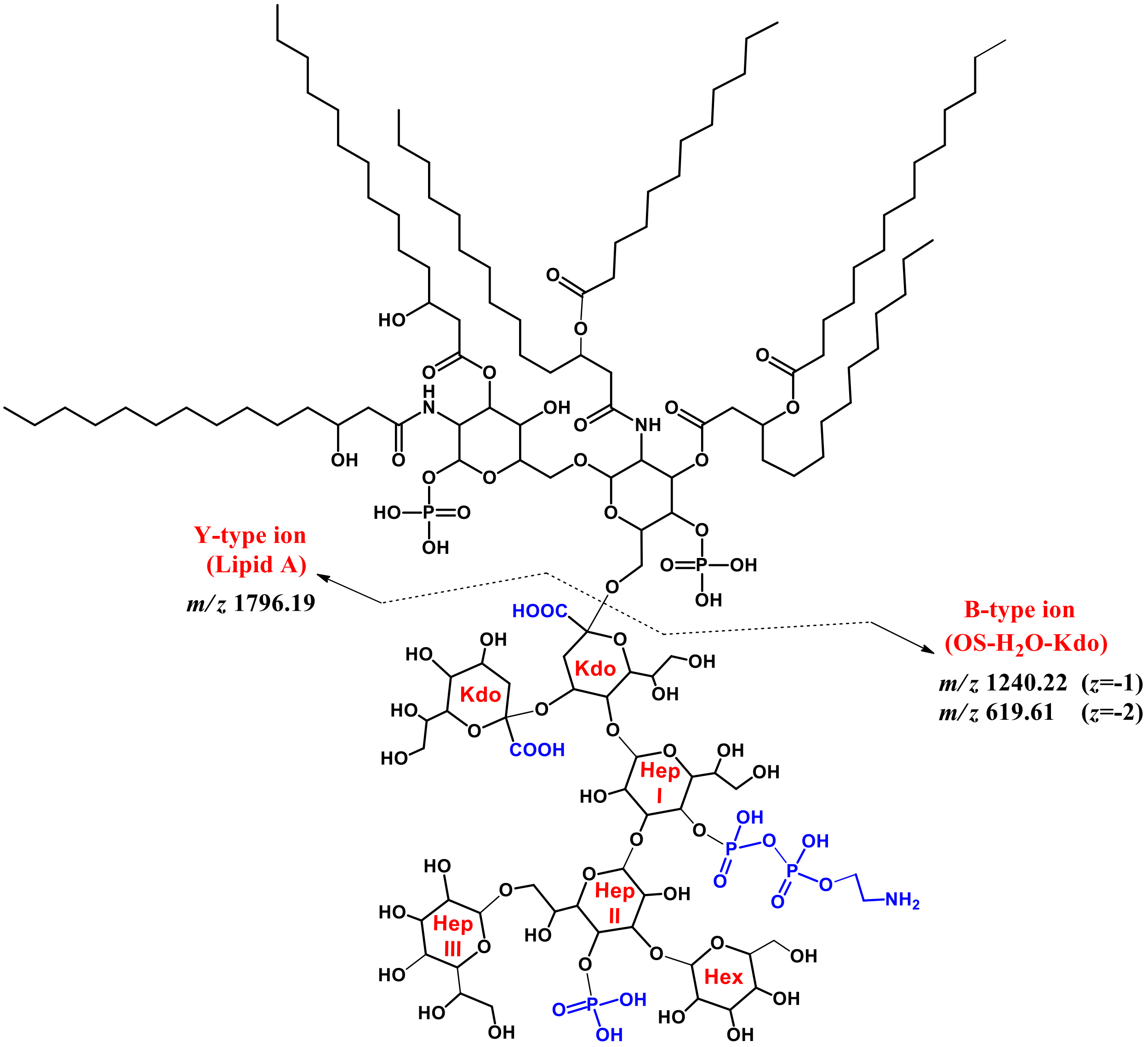


**Figure S5:** Proposed structure of the isobar at *m/z* 1085.17 showing the lipid A and oligosaccharide compositions. It also shows how the Y-type (Lipid A) and B-type (Oligosaccharides) ions/fragments are formed during the source induced dissociation of 100 V.

**Table S1**: List of all product ions originating from the singly charged key ion at *m/z* 993.26. Annotation was created using the GlycoWork bench software and all assignments were within 10 ppm

| Mass to charge | Intensity | Relative Intensity | Ion | Type | Score | Accuracy | Accuracy PPM | Ion m/z | Charges | Neutral Exchanges |
|---|---|---|---|---|---|---|---|---|---|---|
| 993.2628 | 53741.3448 | 62.7753 | 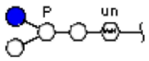 | | 0.0000 | 0.0092 | 9.2443 | 993.2720 | -H | 0 |
| 835.2069 | 53516.4789 | 62.5127 | 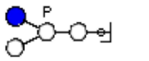 | C | 0.0000 | 0.0057 | 6.7826 | 835.2126 | -H | 0 |
| 817.1986 | 35348.8035 | 41.2910 | 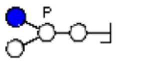 | B | 0.0000 | 0.0034 | 4.1424 | 817.2020 | -H | 0 |
| 813.2030 | 944.6947 | 1.1035 | 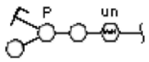 | Z | 0.0000 | 0.0057 | 7.0290 | 813.2087 | -H | 0 |
| 783.1923 | 1047.6561 | 1.2238 | 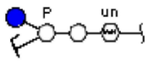 | Z | 0.0000 | 0.0058 | 7.4242 | 783.1981 | -H | 0 |
| 643.1459 | 39730.1194 | 46.4088 | 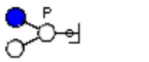 | C | 0.0000 | 0.0033 | 5.1001 | 643.1492 | -H | 0 |
| 643.1459 | 39730.1194 | 46.4088 | 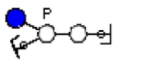 | CY | 0.0000 | 0.0033 | 5.1001 | 643.1492 | -H | 0 |
| 625.1350 | 25702.5073 | 30.0231 | 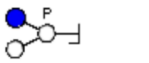 | B | 0.0000 | 0.0036 | 5.8216 | 625.1386 | -H | 0 |
| 625.1350 | 25702.5073 | 30.0231 | 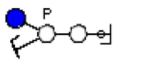 | CZ | 0.0000 | 0.0036 | 5.8216 | 625.1386 | -H | 0 |
| 625.1350 | 25702.5073 | 30.0231 | 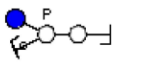 | BY | 0.0000 | 0.0036 | 5.8216 | 625.1386 | -H | 0 |

| Mass to charge | Intensity | Relative Intensity | Ion | Type | Score | Accuracy | Accuracy PPM | Ion m/z | Charges | Neutral Exchanges |
|---|---|---|---|---|---|---|---|---|---|---|
| 607.1246 | 4008.8580 | 4.6828 | 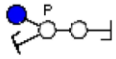  | BZ | 0.0000 | 0.0035 | 5.7654 | 607.1281 | -H | 0 |
| 481.0937 | 959.7565 | 1.1211 | 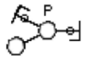  | CY | 0.0000 | 0.0027 | 5.5632 | 481.0964 | -H | 0 |
| 481.0937 | 959.7565 | 1.1211 | 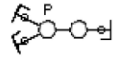  | CYY | 0.0000 | 0.0027 | 5.5632 | 481.0964 | -H | 0 |
| 463.0828 | 9004.8476 | 10.5186 | 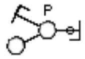  | CZ | 0.0000 | 0.0031 | 6.6023 | 463.0858 | -H | 0 |
| 463.0828 | 9004.8476 | 10.5186 | 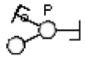  | BY | 0.0000 | 0.0031 | 6.6023 | 463.0858 | -H | 0 |
| 463.0828 | 9004.8476 | 10.5186 | 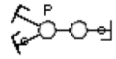  | CYZ | 0.0000 | 0.0031 | 6.6023 | 463.0858 | -H | 0 |
| 463.0828 | 9004.8476 | 10.5186 | 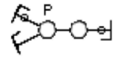  | CZY | 0.0000 | 0.0031 | 6.6023 | 463.0858 | -H | 0 |
| 463.0828 | 9004.8476 | 10.5186 | 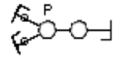  | BYY | 0.0000 | 0.0031 | 6.6023 | 463.0858 | -H | 0 |
| 445.0724 | 9604.0290 | 11.2185 | 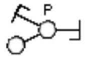  | BZ | 0.0000 | 0.0029 | 6.4587 | 445.0753 | -H | 0 |
| 445.0724 | 9604.0290 | 11.2185 | 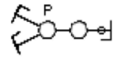  | CZZ | 0.0000 | 0.0029 | 6.4587 | 445.0753 | -H | 0 |
| 445.0724 | 9604.0290 | 11.2185 | 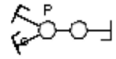  | BYZ | 0.0000 | 0.0029 | 6.4587 | 445.0753 | -H | 0 |
| 445.0724 | 9604.0290 | 11.2185 | 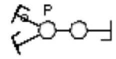  | BZY | 0.0000 | 0.0029 | 6.4587 | 445.0753 | -H | 0 |
| 427.0626 | 2408.4354 | 2.8133 | 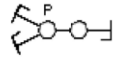  | BZZ | 0.0000 | 0.0021 | 5.0157 | 427.0647 | -H | 0 |

| Mass to charge | Intensity | Relative Intensity | Ion | Type | Score | Accuracy | Accuracy PPM | Ion m/z | Charges | Neutral Exchanges |
|---|---|---|---|---|---|---|---|---|---|---|
| 289.0316 | 2956.8246 | 3.4539 | 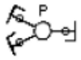 | CYY | 0.0000 | 0.0014 | 4.8562 | 289.0330 | -H | 0 |
| 271.0209 | 13544.2466 | 15.8210 | 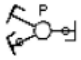 | CYZ | 0.0000 | 0.0015 | 5.5391 | 271.0224 | -H | 0 |
| 271.0209 | 13544.2466 | 15.8210 | 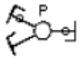 | CZY | 0.0000 | 0.0015 | 5.5391 | 271.0224 | -H | 0 |
| 271.0209 | 13544.2466 | 15.8210 | 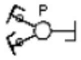 | BYY | 0.0000 | 0.0015 | 5.5391 | 271.0224 | -H | 0 |
| 253.0106 | 11022.3107 | 12.8752 | 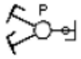 | CZZ | 0.0000 | 0.0013 | 5.0784 | 253.0119 | -H | 0 |
| 253.0106 | 11022.3107 | 12.8752 | 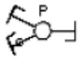 | BYZ | 0.0000 | 0.0013 | 5.0784 | 253.0119 | -H | 0 |
| 253.0106 | 11022.3107 | 12.8752 | 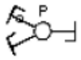 | BZY | 0.0000 | 0.0013 | 5.0784 | 253.0119 | -H | 0 |
| 234.9999 | 4348.7691 | 5.0798 | 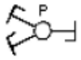 | BZZ | 0.0000 | 0.0014 | 5.9868 | 235.0013 | -H | 0 |

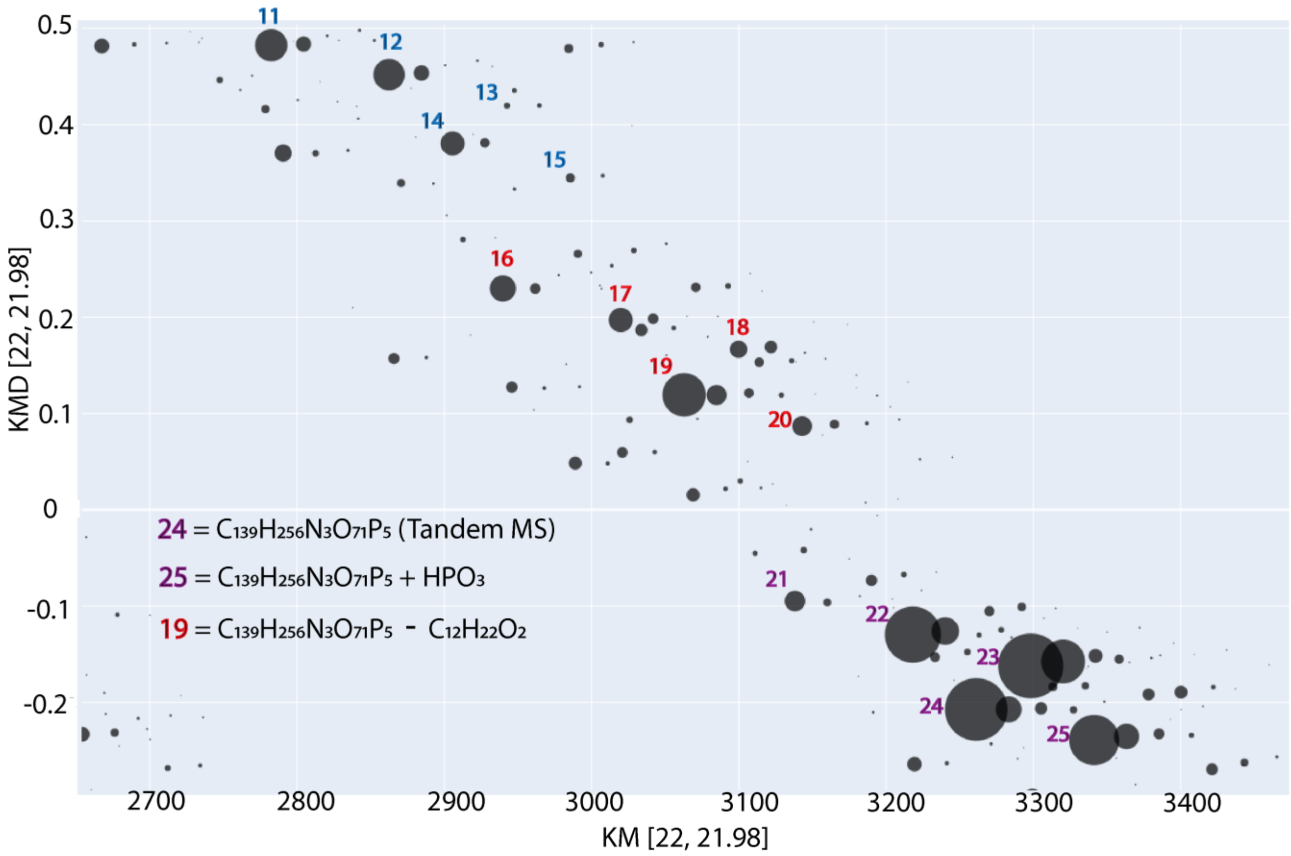


**Figure S6:** KMD plot region showing a constant structural difference of 157.12 Da between ion clusters **11-15 (blue)** and **16-20** (red) and a constant structural difference of 198.16 Da between ion clusters **16-20** (red) and **21-25** (purple). Chemical compositions of ions **24, 25** and **19** are shown on the plot as a representative example for how all the 74 ions compositions were assigned

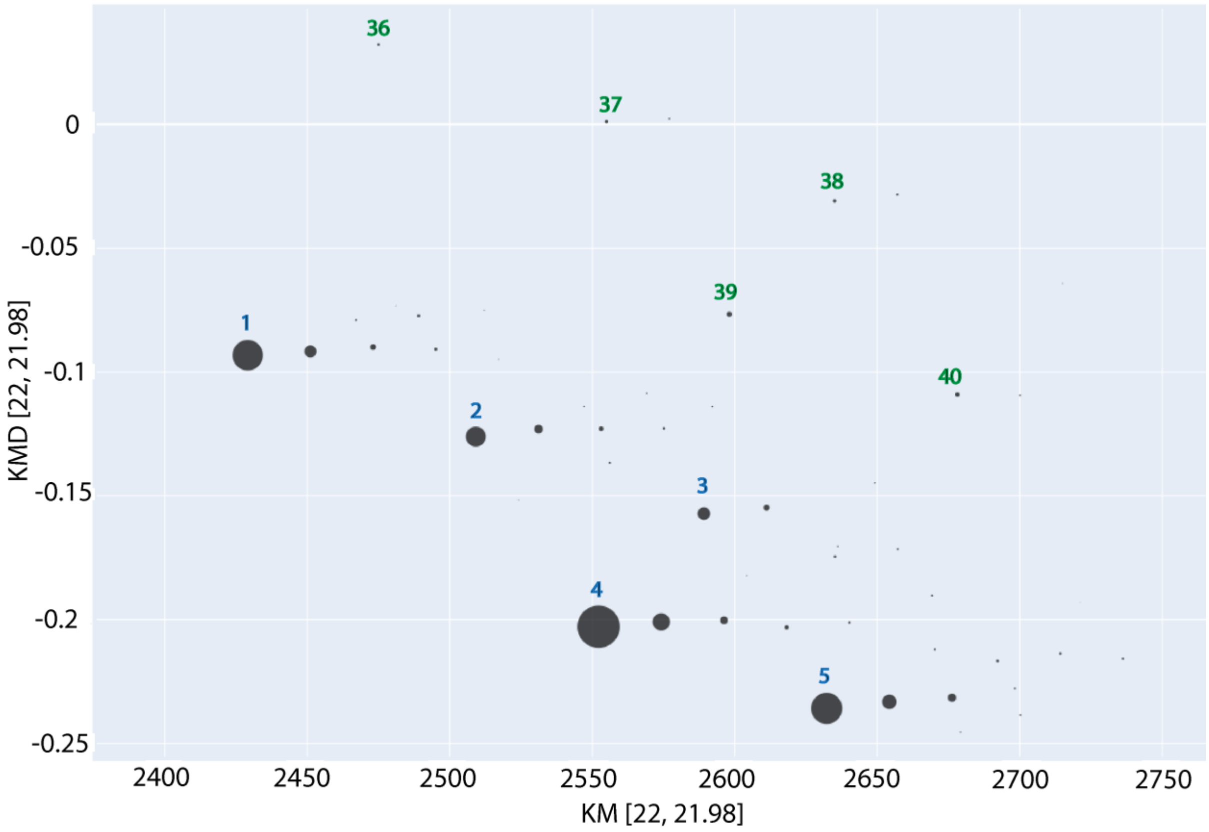


**Figure S7:** KMD plot region showing a constant structural difference of 45.84 Da between ion clusters **1-5** and **36-40** (annotated in blue and green, respectively).

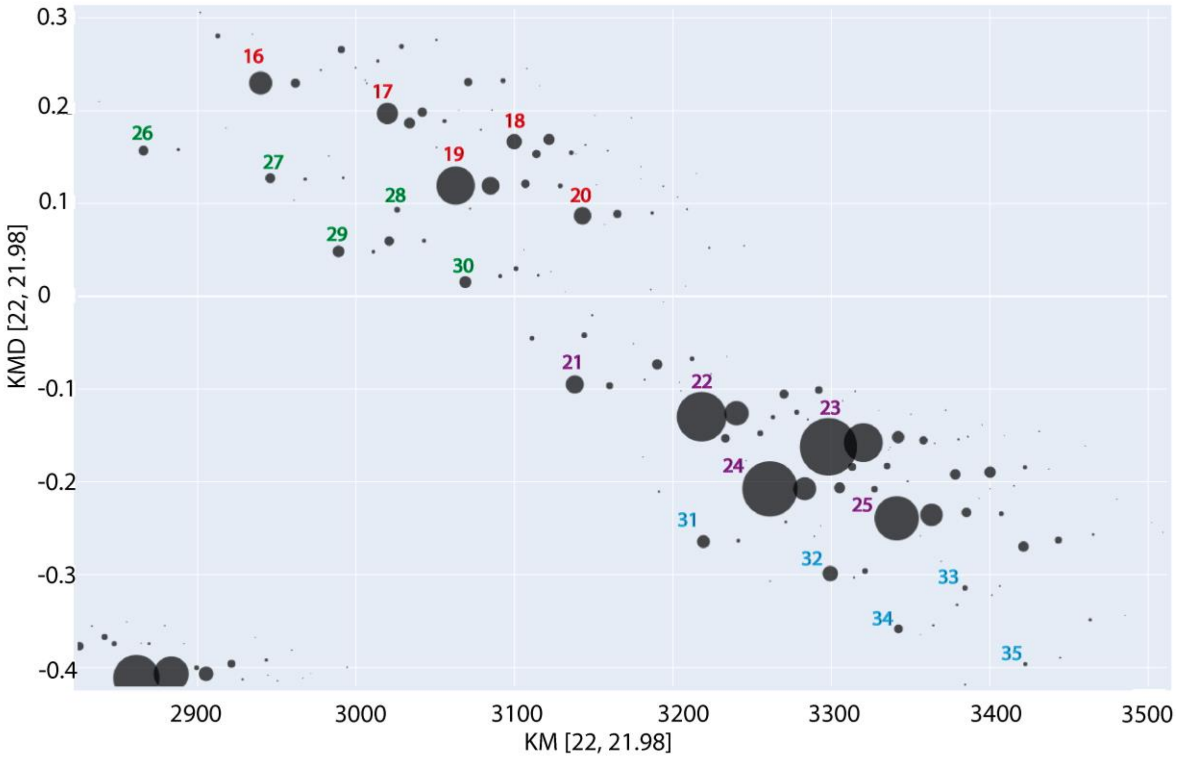


**Figure S8:** KMD plot region showing a constant structural difference of 73.87 Da between ion clusters **16-20** and **26-30** (annotated in red and green, respectively) and a constant structural difference of 81.10 Da between ion clusters **21-25** and **31-35** (annotated in purple and blue, respectively).

**Table S2:** Chemical composition of the *E. coli* J5 LPS (Rc mutant) species revealed form the in-depth KMD plot analysis using the custom base unit of [Na-H]. Abbreviations used are LA: *E. coli* lipid A, Kdo: 3-deoxy- d-manno-oct-2-ulosonic acid, Hep: heptose, Hex: hexose, GlcN: glucosamine, P: phosphate, EN: ethanolamine, 3-hydroxy-tetradecanoic acid: C14:0(3-OH), dodecanoic acid: C12:0, tetradecanoic acid: C14:0

**a) LOS ion species with a complete hexa-acylated lipid A**

| | Molecular weight (Da) | Chemical Composition | Chemical Formula | Error (ppm) | Sodiated Forms |
|---|---|---|---|---|---|
| 1 | 2863.491 | LA+ Kdo2+ Hep2+ Hex+ P | $C_{130}H_{237}N_2O_{59}P_3$ | 3.2 | 2885.4718 |
| 2 | 2943.455 | LA+ Kdo2+ Hep2+ Hex+ P2 | $C_{130}H_{238}N_2O_{62}P_4$ | 2.3 | 2965.438 |
| 3 | 2986.4986 | LA+ Kdo2+ Hep2+ Hex+ P2+ EN | $C_{132}H_{243}N_3O_{62}P_4$ | 2.7 | 3008.4809 |
| 4 | 3023.4233 | LA+ Kdo2+ Hep2+ Hex+ P3 | $C_{130}H_{239}N_2O_{65}P_5$ | 2.9 | 3045.4008 |
| 5 | 3066.4659 | LA+ Kdo2+ Hep2+ Hex+ P3+ EN | $C_{132}H_{244}N_3O_{65}P_5$ | 3.0 | 3088.4415 |
| 6 | 3108.492 | LA+ Kdo2+ Hep2+ Hex+ P3+ EN+ [$C_3H_6$] | $C_{135}H_{250}N_3O_{65}P_5$ | -3.8 | |
| 7 | 3135.5195 | LA + Kdo2+ Hep3+ Hex+ P2 | $C_{137}H_{250}N_2O_{68}P_4$ | 2.5 | 3157.5029, 3179.4783 |
| 8 | 3146.436 | LA+ Kdo2+ Hep2+ Hex+ P4+ EN | $C_{132}H_{245}N_3O_{68}P_6$ | 4.08 | |
| 9 | 3187.4553 | LA (-$C_2H_4$) + Kdo2+ Hep3+Hex+P3 | $C_{135}H_{247}N_2O_{71}P_5$ | 2.7 | 3209.4313, 3231.4107 |
| 10 | 3215.4886 | LA + Kdo2+ Hep3+ Hex+ P3 | $C_{137}H_{251}N_2O_{71}P_5$ | 3.3 | 3237.4669 |
| 11 | 3216.6223 | LA+ Kdo2+ Hep3+ Hex+ GlcN+ P | $C_{143}H_{260}N_3O_{69}P_3$ | 2.5 | 3238.6032 |
| 12 | 3258.531 | LA + Kdo2+ Hep3+ Hex+ P3+ EN | $C_{139}H_{256}N_3O_{71}P_5$ | 3.3 | 3280.5129, 3302.4938, 3324.4772 |
| 13 | 3267.4216 | LA (-$C_2H_4$) + Kdo2+ Hep3+Hex+P4 | $C_{135}H_{248}N_2O_{74}P_6$ | 2.6 | 3289.399 |
| 14 | 3295.4554 | LA + Kdo2+ Hep3+ Hex+ P4 | $C_{137}H_{252}N_2O_{74}P_6$ | 3.4 | 3317.4328, 3339.4089 |
| 15 | 3296.5911 | LA+ Kdo2+ Hep3+ Hex+ GlcN+ P2 | $C_{143}H_{261}N_3O_{72}P_4$ | 3.2 | 3318.5703 |
| 16 | 3338.4971 | LA + Kdo2+ Hep3+ Hex+ P4+ EN | $C_{139}H_{257}N_3O_{74}P_6$ | 3.2 | 3360.4753, 3382.4546, 3404.438 |

| | | | | | |
|---|---|---|---|---|---|
| 17 | 3339.6153 | LA+ Kdo2+ Hep3+ Hex+ GlcN+ P2+ EN | $C_{145}H_{266}N_4O_{72}P_4$ | -2.2 | 3361.5935 |
| 18 | 3375.4194 | LA + Kdo2+ Hep3+ Hex+ P5 | $C_{137}H_{253}N_2O_{77}P_7$ | 2.6 | 3397.399, 3419.3757 |
| 19 | 3381.5368 | LA + Kdo2+ Hep3+ Hex+ P4+ EN2 | $C_{141}H_{262}N_4O_{74}P_6$ | 2.4 | 3403.5167 |
| 20 | 3381.6409 | LA+ Kdo2+ Hep3+ Hex+ GlcN+ P2+ EN+ Ac | $C_{147}H_{268}N_4O_{73}P_4$ | 2.3 | |
| 21 | 3418.4618 | LA + Kdo2+ Hep3+ Hex+ P5+ EN | $C_{139}H_{258}N_3O_{77}P_7$ | 2.6 | 3440.4367, 3462.4128 |
| 22 | 3419.5876 | LA+ Kdo2+ Hep3+ Hex+ GlcN+ P3+ EN | $C_{145}H_{267}N_4O_{75}P_5$ | -0.4 | 3441.5628 |
| 23 | 3461.6094 | LA+ Kdo2+ Hep3+ Hex+ GlcN+ P3+ EN+ Ac | $C_{147}H_{269}N_4O_{76}P_5$ | 2.8 | |

**b) LOS ion species with penta-acylated lipid A**

| | Molecular weight (Da) | Chemical Composition | Chemical Formula | Error (ppm) | Sodiated forms |
|---|---|---|---|---|---|
| 1 | 2665.3291 | (LA - C12:0 - O) + Kdo2+ Hep2+ Hex+ P | $C_{118}H_{215}N_2O_{57}P_3$ | 3.4 | 2687.3094, 2709.2899 |
| 2 | 2745.2986 | (LA - C12:0 - O) + Kdo2+ Hep2+ Hex+ P2 | $C_{118}H_{216}N_2O_{60}P_4$ | 4.5 | 2767.2762 |
| 3 | 2776.3035 | (LA - C14:0) + Kdo2+ Hep2+ Hex+ P2+ EN | $C_{118}H_{217}N_3O_{61}P_4$ | 4.1 | 2798.2761 |
| 4 | 2788.3393 | (LA - C12:0 - O) + Kdo2+ Hep2+ Hex+ P2+ EN | $C_{120}H_{221}N_3O_{60}P_4$ | 3.9 | 2810.3214, 2832.3003 |
| 5 | 2868.3048 | (LA - C12:0 - O) + Kdo2+ Hep2+ Hex+ P3+ EN | $C_{120}H_{222}N_3O_{63}P_5$ | 3.5 | 2890.2872 |
| 6 | 2910.3289 | (LA - C12:0 - O) + Kdo2+ Hep2+ Hex+ P3+ EN+ [$C_3H_6$] | $C_{123}H_{228}N_3O_{63}P_5$ | -4.4 | 2932.3091 |
| 7 | 2937.3575 | (LA - C12:0 - O) + Kdo2+ Hep3+ Hex+ P2 | $C_{125}H_{228}N_2O_{66}P_4$ | 2.6 | 2959.3396 |
| 8 | 3017.3246 | (LA - C12:0 - O) + Kdo2+ Hep3+ Hex+ P3 | $C_{125}H_{229}N_2O_{69}P_5$ | 2.8 | 3039.3052 |
| 9 | 3018.4612 | (LA - C12:0 - O) + Kdo2+ Hep3+ Hex+ GlcN+ P | $C_{131}H_{238}N_3O_{67}P_3$ | 3.0 | 3040.4428 |
| 10 | 3060.367 | (LA - C12:0 - O) + Kdo2+ Hep3+ Hex+ P3+ EN | $C_{127}H_{234}N_3O_{69}P_5$ | 2.9 | 3082.3491, 3104.329, 3126.3131 |
| 11 | 3097.2893 | (LA - C12:0 - O) + Kdo2+ Hep3+ Hex+ P4 | $C_{125}H_{230}N_2O_{72}P_6$ | 2.2 | 3119.2689 |
| 12 | 3098.4252 | (LA - C12:0 - O) + Kdo2+ Hep3+ Hex+ GlcN+ P2 | $C_{131}H_{239}N_3O_{70}P_4$ | 2.2 | 3120.4102 |

| | | | | | |
|---|---|---|---|---|---|
| 13 | 3140.3338 | (LA - C12:0 - O) + Kdo2+ Hep3+ Hex+ P4+ EN | $C_{127}H_{235}N_{3}O_{72}P_{6}$ | 2.9 | 3162.3138, 3184.2947, 3206.2727 |
| 14 | 3141.4617 | (LA - C12:0 - O) + Kdo2+ Hep3+ Hex+ GlcN+ P2+ EN | $C_{133}H_{244}N_{4}O_{70}P_{4}$ | 0.3 | |
| 15 | 3220.3026 | (LA - C12:0 - O) + Kdo2+ Hep3+ Hex+ P5+ EN | $C_{127}H_{236}N_{3}O_{75}P_{7}$ | 3.6 | 3242.2824 |

**c) LOS ion species with tetra-acylated lipid A**

| | Molecular weight (Da) | Chemical Composition | Chemical Formula | Error (ppm) | Sodiated forms |
|---|---|---|---|---|---|
| 1 | 2427.0997 | (LA - C14:0(3-OH) - C14:0)+ Kdo2+ Hep2+ Hex+ P | $C_{102}H_{185}N_{2}O_{56}P_{3}$ | 3.9 | 2449.0801, 2471.0603, 2493.0431 |
| 2 | 2507.0669 | (LA - C14:0(3-OH) - C14:0)+ Kdo2+ Hep2+ Hex+ P2 | $C_{102}H_{186}N_{2}O_{59}P_{4}$ | 4.1 | 2529.0458, 2551.0276, 2573.0094 |
| 3 | 2550.1083 | (LA - C14:0(3-OH) - C14:0)+ Kdo2+ Hep2+ Hex+ P2+ EN | $C_{104}H_{191}N_{3}O_{59}P_{4}$ | 3.7 | 2572.0883, 2594.0696, 2616.0544, 2638.0344 |
| 4 | 2587.0323 | (LA - C14:0(3-OH) - C14:0) + Kdo2+ Hep2+ Hex+ P3 | $C_{102}H_{187}N_{2}O_{62}P_{5}$ | 3.6 | 2609.0118 |
| 5 | 2630.0756 | (LA - C14:0(3-OH) - C14:0) + Kdo2+ Hep2+ Hex+ P3+ EN | $C_{104}H_{192}N_{3}O_{62}P_{5}$ | 4.0 | 2652.0549, 2674.0352, 2696.0134, 2717.9967 |
| 6 | 2666.9997 | (LA - C14:0(3-OH) - C14:0) + Kdo2+ Hep2+ Hex+ P4 | $C_{102}H_{188}N_{2}O_{65}P_{6}$ | 3.9 | |
| 7 | 2673.1167 | (LA - C14:0(3-OH) - C14:0) + Kdo2+ Hep2+ Hex+ P3+ EN2 | $C_{106}H_{197}N_{4}O_{62}P_{5}$ | 3.5 | 2695.0965, 2717.0761 |
| 8 | 2699.1303 | (LA - C14:0(3-OH) - C14:0)+ Kdo2+ Hep3+ Hex+ P2 | $C_{109}H_{198}N_{2}O_{65}P_{4}$ | 3.8 | 2721.1101, 2743.0912, 2765.0741, 2787.0531 |
| 9 | 2710.0424 | (LA- C14:0(3-OH) - C14:0) + Kdo2+ Hep2+ Hex+ P4+ EN | $C_{104}H_{193}N_{3}O_{65}P_{6}$ | 4.0 | 2732.0217 |
| 10 | 2753.083 | (LA - C14:0(3-OH) - C14:0)+ Kdo2+ Hep2+ Hex+ P4+ EN2 | $C_{106}H_{198}N_{4}O_{65}P_{6}$ | 3.3 | 2775.0616, 2797.0381 |
| 11 | 2779.0984 | (LA - C14:0(3-OH) - C14:0)+ Kdo2+ Hep3+ Hex+ P3 | $C_{109}H_{199}N_{2}O_{68}P_{5}$ | 4.3 | 2801.077, 2823.0583, 2845.0376, |

| | | | | | |
|---|---|---|---|---|---|
| | | | | | 2867.0194 |
| 12 | 2780.2339 | (LA - C14:0(3-OH) - C14:0) + Kdo2+ Hep3+ Hex+ GlcN+ P | $C_{115}H_{208}N_3O_{66}P_3$ | 4.1 | 2802.2146 |
| 13 | 2822.1404 | (LA - C14:0(3-OH) - C14:0)+ Kdo2+ Hep3+ Hex+ P3+ EN | $C_{111}H_{204}N_3O_{68}P_5$ | 4.2 | 2844.1206, 2866.1015, 2888.0858 |
| 14 | 2859.063 | (LA - C14:0(3-OH) - C14:0)+ Kdo2+ Hep3+ Hex+ P4 | $C_{109}H_{200}N_2O_{71}P_6$ | 3.6 | 2881.0406, 2903.0224 |
| 15 | 2860.1988 | (LA - C14:0(3-OH) - C14:0)+ Kdo2+ Hep3+ Hex+ GlcN+ P2 | $C_{115}H_{209}N_3O_{69}P_4$ | 3.5 | 2882.179 |
| 16 | 2902.1054 | (LA - C14:0(3-OH) - C14:0)+ Kdo2+ Hep3+ Hex+ P4+ EN | $C_{111}H_{205}N_3O_{71}P_6$ | 3.6 | 2924.084, 2946.0642, 2968.0464, 2990.028, 3012.0138, 3033.9932 |
| 17 | 2903.2349 | (LA - C14:0(3-OH) - C14:0)+ Kdo2+ Hep3+ Hex+ GlcN+ P2+ EN | $C_{117}H_{214}N_4O_{69}P_4$ | 1.3 | 2925.2162 |
| 18 | 2939.028 | (LA - C14:0(3-OH) - C14:0) + Kdo2+ Hep3+ Hex+ P5 | $C_{109}H_{201}N_2O_{74}P_7$ | 3.0 | 2961.0041 |
| 19 | 2940.1653 | (LA - C14:0(3-OH) - C14:0) +Kdo2+ Hep3+ Hex+ GlcN+ P3 | $C_{115}H_{210}N_3O_{72}P_5$ | 3.4 | 2962.1471 |
| 20 | 2945.2477 | (LA - C14:0(3-OH) - C14:0) + Kdo2+ Hep3+ Hex+ GlcN+ P2+ EN+ AC | $C_{119}H_{216}N_4O_{70}P_4$ | 2.1 | |
| 21 | 2982.0715 | (LA - C14:0(3-OH) - C14:0) + Kdo2+ Hep3+ Hex+ P5+ EN | $C_{111}H_{206}N_3O_{74}P_7$ | 3.4 | 3004.0495, 3026.0287 |
| 22 | 2983.205 | (LA - C14:0(3-OH) - C14:0) + Kdo2+ Hep3+ Hex+ GlcN+ P3+ EN | $C_{117}H_{215}N_4O_{72}P_5$ | 2.5 | 3005.1845 |
| 23 | 2988.2797 | (LA - C14:0(3-OH) - C14:0) + Kdo2+ Hep3+ Hex+ GlcN+ P2+ EN2+ AC | $C_{121}H_{221}N_5O_{70}P_4$ | -1.4 | |
| 24 | 3026.245 | (LA - C14:0(3-OH) - C14:0)+ Kdo2+ Hep3+ Hex+ GlcN+ P3+ EN2 | $C_{119}H_{220}N_5O_{72}P_5$ | 1.8 | 3048.2200 |
| 25 | 3031.3235 | (LA - C14:0(3-OH) - C14:0) + Kdo2+ Hep3+ Hex+ GlcN+ P2+ EN3+ AC | $C_{123}H_{226}N_6O_{70}P_4$ | -0.8 | 3053.3033 |
| 26 | 3068.2489 | (LA - C14:0(3-OH) - C14:0) + Kdo2+ Hep3+ Hex+ GlcN+ P3+ EN2+ AC | $C_{121}H_{222}N_5O_{73}P_5$ | -0.4 | 3090.2294 |
| 27 | 3111.2912 | (LA - C14:0(3-OH) - C14:0)+ Kdo2+ Hep3+ Hex+ GlcN+ P3+ EN3+ AC | $C_{123}H_{227}N_6O_{73}P_5$ | -0.36 | 3133.2716 |

**d) LOS ion species with tri-acylated lipid A**

| | Molecular weight (Da) | Chemical Composition | Chemical Formula | Error (ppm) | Sodiated forms |
|---|---|---|---|---|---|
| 1 | 2323.9143 | (LA - 2 x C14:0(3-OH) - C14:0) + Kdo2+ Hep2+ Hex+ P2+ EN | $C_{90}H_{165}N_3O_{57}P_4$ | 3.8 | 2345.8944 |
| 2 | 2367.9416 | (LA - C14:0(3-OH) -C14:0 - C12:0) + Kdo2+ Hep2+ Hex+ P2+ EN | $C_{92}H_{169}N_3O_{58}P_4$ | 4.2 | 2389.9217 |
| 3 | 2403.8815 | (LA - 2 x C14:0(3-OH) - C14:0) + Kdo2+ Hep2+ Hex+ P3+ EN | $C_{90}H_{166}N_3O_{60}P_5$ | 4.0 | 2425.8609 |
| 4 | 2472.9366 | (LA - 2 x C14:0(3-OH) - C14:0)+ Kdo2+ Hep3+ Hex+ P2 | $C_{95}H_{172}N_2O_{63}P_4$ | 4.0 | |
| 5 | 2552.902 | (LA - 2 x C14:0(3-OH) - C14:0)+ Kdo2+ Hep3+ Hex+ P3 | $C_{95}H_{173}N_2O_{66}P_5$ | 3.5 | 2574.8828 |
| 6 | 2595.9445 | (LA - 2 x C14:0(3-OH) - C14:0)+ Kdo2+ Hep3+ Hex+ P3+ EN | $C_{97}H_{178}N_3O_{66}P_5$ | 3.5 | 2617.9248 |
| 7 | 2632.8684 | (LA - 2 x C14:0(3-OH) - C14:0)+ Kdo2+ Hep3+ Hex+ P4 | $C_{95}H_{174}N_2O_{69}P_6$ | 3.4 | 2654.8478 |
| 8 | 2675.9112 | (LA - 2 x C14:0(3-OH) - C14:0)+ Kdo2+ Hep3+ Hex+ P4+ EN | $C_{97}H_{179}N_3O_{69}P_6$ | 3.6 | 2697.8935 |
